# Preferential Isoform Usage and Isoform Switching in Systemic Lupus Erythematosus Patients with Distinct Autoantibody Profile

**DOI:** 10.64898/2026.11.05.764679

**Authors:** Richa Rai, Arya Das, Sudhir Kumar Chauhan, Vikas Vikram Singh, Madhukar Rai, Dinesh Velayutham, Geeta Rai

## Abstract

RNA sequencing has advanced our understanding of molecular heterogeneity in systemic lupus erythematosus (SLE), yet most studies have focused on differential gene expression (DGE), which overlooks transcript-level complexity. Alternative splicing and polyadenylation generate multiple isoforms from a single gene, and their dysregulation may contribute to autoimmune pathogenesis, including through the production of non-canonical isoforms with potential autoantigenic properties. To explore this underexamined dimension, we analysed differential isoform usage (DIU) and major isoform switching (MIS) across SLE subgroups stratified by autoantibody profile: anti-dsDNA^+^, anti-ENA^+^ and anti-dsDNA^+^ENA^+^. Subgroup-specific DIU and MIS events were enriched in RNA-binding and transcription-related functions. In the anti-dsDNA^+^ subgroup, the T-cell receptor adaptor GRAP2 showed isoform switching accompanied by expansion of a non-coding retained-intron transcript. In the anti-ENA^+^ subgroup, IGHM isoform switched from the membrane bound to the secretory form, consistent with an antibody secreting cell signature; NACA, a reported IgE autoantigen, showed isoform switching; and BRI3, linked to TNF-induced cell death, switched toward its protein-coding transcript. In the anti-dsDNA^+^ENA^+^ subgroup, TNFAIP2, a TNF-induced gene, showed isoform redistribution with loss of a uORF containing transcript. These findings highlight isoform regulation as a distinct, subgroup-specific and functionally relevant layer of transcriptomic dysregulation in SLE.

## Introduction

RNA-sequencing has significantly enhanced our understanding in characterizing molecular and functional landscape pertaining to distinct clinical phenotypes in systemic lupus erythematosus (SLE) patients. Most analyses to date are predominantly focused on the identification of differentially expressed genes (DEG) which scores the aggregated expression of all the isoforms (or transcripts) originating from a gene [1–4]. These DEGs were significantly enriched in immune-related pathways, which offered critical insights into the SLE pathogenesis [1, 2, 5–7]. However, it fails to capture the dynamics in the expression of multiple related isoforms [8].

Alternative splicing generates multiple isoforms from a single gene, where one isoform predominates over others, this is often termed the “canonical” or “dominant” isoforms. Although alternative splicing is a normal regulatory mechanism, disruption of normal splicing pattern can have detrimental consequences on disease pathology [9–12]. In autoimmune disease, aberrant splicing events may lead to the production of non-canonical isoforms that could contribute to the generation of isoforms which could serve as an autoantigen [13–15]. Nevertheless, alternative splicing such as exon skipping, intron retention, or alternative 5’or 3’ splice site usage remains poorly characterized in SLE and warrant further investigation.

Isoform level analysis, referred to as differential isoform usage (DIU) addresses this gap by identifying changes in isoform proportion and major isoform switching (MIS), together with their predicted functional consequences, which may be overlooked in classical DEG analyses [16]. Notably, DIU genes have not been systematically studied in SLE or other autoimmune diseases. Although, we have previously identified differential gene and transcript expression in SLE [1], gene level analysis lacks the resolution to detect transcript level changes such as alternative splicing or isoform switching and their functional consequences. We therefore analyzed DIU and MIS across SLE subgroups defined by autoantibody profiles to determine their association with disease heterogeneity.

Many genes with minimal expression changes (logFC < 2) showed substantial shifts in isoform usage (up to 73%), indicating that isoform level alterations can occur independently of gene level changes. Subgroup specific DIU and MIS events were enriched in RNA-binding and transcription related functions. In the anti-dsDNA^+^ subgroup, the T-cell receptor adaptor GRAP2 showed isoform switching accompanied by expansion of a non-coding retained-intron transcript. In the anti-ENA^+^ subgroup, IGHM isoform switched from the membrane-bound to the secretory form; NACA, a reported IgE autoantigen, showed isoform switching; and BRI3, linked to TNF-α-induced cell death, switched toward its protein-coding transcript. In the anti-dsDNA^+^ENA^+^ subgroup, the TNF-α-induced gene TNFAIP2 showed isoform redistribution with loss of a (upstream open reading frame) uORF containing transcript. These findings suggest that isoform regulation constitutes a distinct, subgroup-specific layer of transcriptomic dysregulation in SLE.

## Methodology

### Dataset

We used our previously published dataset (GEO: GSE80183), which involves 12 SLE patients and 4 controls [1]. SLE patients were segregated based on distinct autoantibodies that are directed against either (i) dsDNA or (ii) RNA-associated proteins (also known as extractable nuclear antigens, ENA) or (iii) both (dsDNA and ENA). Earlier, this sub-grouping approach revealed interesting molecular phenotypes associated with each SLE subsets, like differential expression of toll-like receptors (TLRs) [6], heat shock proteins (HSPs) [7], miRNA-based genetic regulation [17], divergent sources of autoantigens resulting from defective clearance mechanisms, either phagocytosis or NETosis or both [18] and specific transcriptome profiles [1].

### Differential Isoform Usage and Isoform Switching Events

TappAS computation tool was used to investigate isoform switching and alternative splicing in different SLE patients’ subsets [19]. Briefly, the reads obtained from Illumina Hiseq 2500 were filtered and mapped to hg38 genome, to determine the abundance of transcripts using Cuffdiff [20, 21]. The transcript expression matrix and transcript-level functional annotation, along with the experimental design matrix were uploaded to the TappAS tool. This tool uses data from multiple functional databases and is integrated with the transcript-level expression data to create isoform-level annotations (of domains, motifs, and sites) for both coding and non-coding isoforms [19]. This tool gives information on isoform switching events as MIS.

#### Major isoform switch

The major isoform often refers to the isoform of a gene which has the highest mean expression whereas the remaining isoforms are defined as minor isoforms in specific experimental conditions. The term “major” only provides a reference point and does not necessarily reflect the functional relevance whereas isoform switching in the given condition may reflect implications in pathological conditions. Isoform switch is defined as an event where the relative contribution of the isoforms to the respective gene expression changes significantly between experimental/ diseased conditions **(Supplementary Figure 1)**. We determined the isoform switch by calculating the isoform fraction (IF) which is the ratio of Fragments Per Kilobase of transcript per Million mapped reads (FPKM) of each isoform to the total gene expression. Then, the difference in the IF under experimental/ diseased condition is calculated as differential isoform fraction (dIF) [dIF = IF _(Diseased Group)_-IF _(Control)_]. Isoform switch is considered as a significant event if mean dIF > 0.1 and FDR <0.05 [22, 23]. Switching events identify differential feature inclusion, differential polyadenylation, and differential isoform usage events that change direction between conditions and can be used to prioritize candidates for further analysis.

#### Differential isoform usage

DIU is another term that is concurrently used to show the isoform switch. However, it’s not same, DIU is defined by the magnitude of difference in expression of isoform in a gene across given diseased conditions which is considered significant or not based on the threshold of significance given (Default threshold is 0.05). This tool analyses DIU by testing multiple isoforms in a single model to determine total usage change (TUC) and quantify the magnitude of changes for DIU genes. TUC measures the amount of redistribution (as %) in the expression levels across different conditions for isoforms of the same gene. A typical cut-off for TUC is > 50%. This means that if the relative expression of one isoform compared to other changes by more than 50% between two conditions (e.g., SLE patients vs controls), it is considered a significant change. At some instances, more conservative cut-off values like 20% or 30% for total usage change is employed to look for more subtle shifts in isoform usage to see their biologically significance. The log2 fold change (logFC) was also used in combination with TUC to assess DIU. For instance, if an isoform shows a log2FC>1 (which corresponds to a fold change of 2), and the TUC is > 50%, this will indicate a strong and biologically significant switch in isoform usage.

#### Functional feature inclusion (FI) and functional diversity analysis (FDA)

Differential feature inclusion (DFI) reflects the changes in the molecular features (such as exons, UTRs, miRNA binding sites, uORFs, etc.) between different isoforms of a gene. In the context of SLE or other diseases, this can have profound effects on gene regulation, protein function, and disease mechanisms. This is a qualitative analysis of functional diversity in a transcriptome annotation. It measures the changes in the isoforms of a gene by interrogating functional features like 3’- and 5’-UTR length, CDS length, 3’- and 5’-UTR motifs, poly adenylation site (PAS), uORF and miRNA binding site that vary across isoforms from the same gene. If the changes in any of the mentioned transcript or protein annotated sites are observed the status is represented as “varying” whereas no change is represented as “not varying”. This tool assesses the varying status of individual genes for each functional feature category, and by feature ID/ annotation, i.e., assessing the number of genes for which a particular feature is differentially present across isoforms. Varying status is computed both by genomic position (positional approach) and by the presence/absence (presence approach) - depending on the type of feature under evaluation. Additionally, isoform usage is considered “varying” if the disease condition leads to significant changes in splicing patterns, which may impact gene function and contribute to disease mechanisms. If there is no significant change in isoform expression between the disease and control states, isoform usage is considered “not varying”, implying that the gene may not be directly involved in the disease through alternative splicing mechanisms.

Ensemble genome browser GRCh38 was used to retrieve the details corresponding to transcript ID of each gene including transcript size, protein size, isoform biotype, CCDS, Uniprot Match and RefSeq Match. Further, the information based on usage change from most dominant isoform to any of the biotype whether its protein coding, processed transcript, retained intron it was determined if it impacts the functional property of a gene in diseased condition.

## Results

### Isoform usage change irrespective of differential gene expression

We investigated the isoform switching and usage change to determine if the change in these events associates with specific SLE subgroups which were classified based on distinct autoantibody repertoire. These molecular events could majorly contribute to the pathology of the disease which could be missed if focused only on the gene expression. Therefore, we sought to compare if these events are independent of the DEG. In all the SLE patients as a single group as well as individual subsets, we observed that majority of the genes that show differential expression of logFC between 2, they also represent the total usage change of more than 50% indicating that switching events and total usage change occurs regardless of gene expression change (**Figure 1 A-D**). Comparing all SLE patients as a single group with healthy individual resulted in 959 genes that show MIS and six genes shows DIU. However, when we segregated SLE patients into three subsets we observed 776 MIS and ten DIU in anti-dsDNA^+^ subgroup; 782 MIS and fifteen DIU in anti-ENA^+^ subgroup; and 834 MIS and seven DIU genes in anti-dsDNA^+^ENA^+^ subgroup (**Supplementary figure 2A**). Furthermore, we performed the gene ontology (GO) enrichment analysis of all the MIS genes using the DAVID bioinformatic resource [24] for molecular functions of the uniquely observed MIS genes in each SLE subsets. We found that these genes are enriched in protein binding, RNA binding, RNA polymerase II promoter binding, Ubiquitin like protein binding functions in either one or more SLE patient subsets (**Supplementary table 1 and Supplementary figure 2B and C**). Interestingly, the DIU genes have not been studied in SLE or autoimmune disease in much detail therefore, in depth analysis of DIU genes was performed. Among the identified DIU genes there are only few genes which also demonstrates major isoform switching with dIF > 0.1 (**Supplementary table 2**).

**Figure 1.**
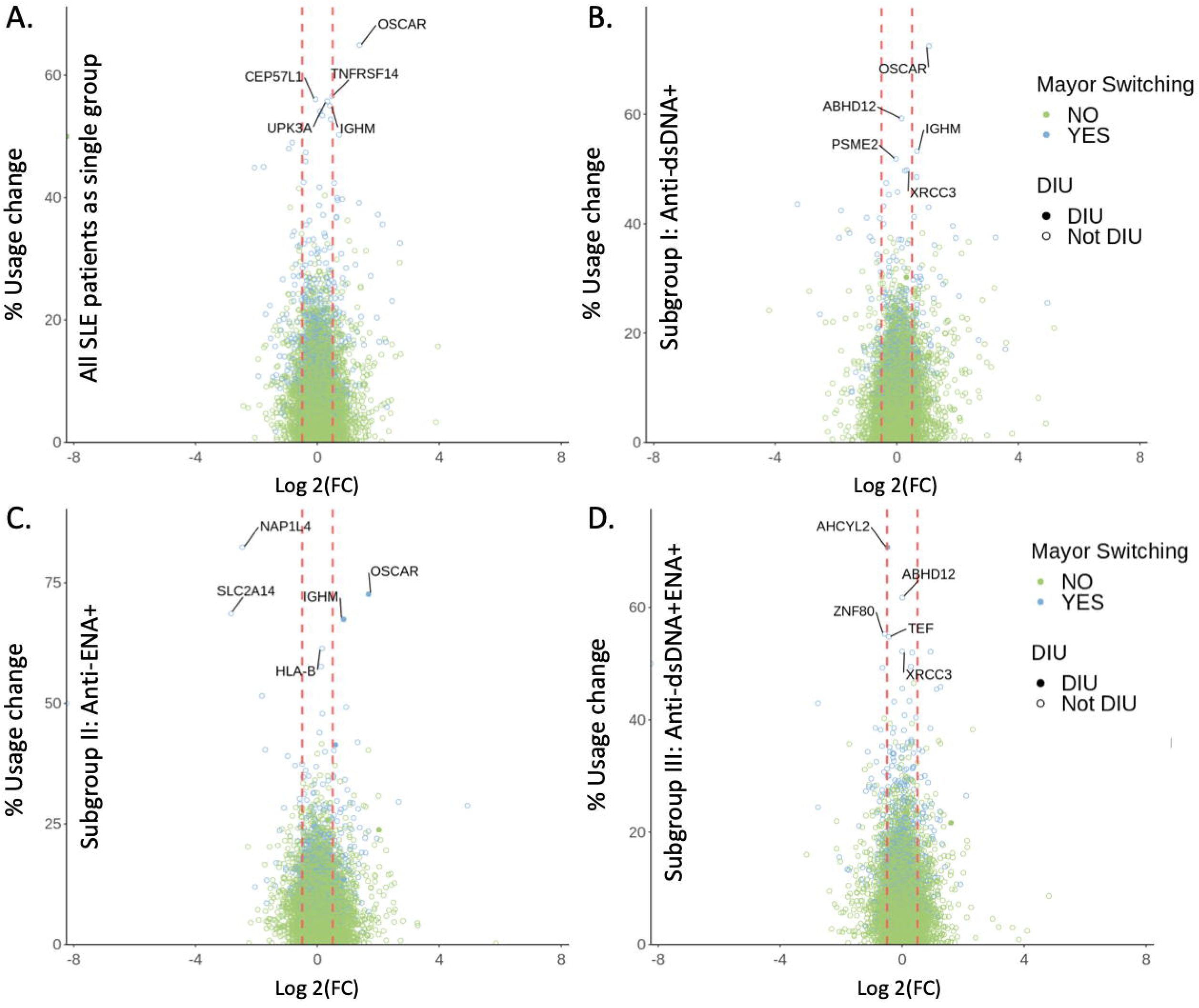
Graphical representation of the genes showing DIU and MIS. **A.** all the SLE patients as a single group **B.** Anti-dsDNA^+^ subset **C.** Anti-ENA^+^ subset **D.** Anti-dsDNA^+^ENA^+^ subsets. Blue open circle represents MIS only whereas blue closed circle represents genes that are both MIS and DIU.

### Isoform usage change and isoform switching events predominant in anti-ENA^+^ patients

We identified distinct sets of DIU genes in each SLE patients’ subsets, largely involving non-overlapping gene sets. These includes ten DIU genes (DAZAP2, FAM96A, FAU, GRAP2, HCK, NAA38, SLC2A3, TNFAIP2, YPEL3 and ZSWIM8) in anti-dsDNA^+^ patients out of which seven (DAZAP2, FAU, GRAP2, NAA38, SLC2A3, YPEL3 and ZSWIM8) genes were unique in this group. 13 out of 15 DIU genes were unique to anti-ENA^+^ subgroups which includes ADORA2A, AP2M1, BRI3, CRIP1, IGHM, ITGAL, NACA, OSCAR, PTBP1, RALB, RPL41, SAMD3 and TMBIM6 whereas anti-dsDNA^+^ENA^+^ subgroup shows only three unique DIU, ADIPOR1, AHCYL2 and IFT20 **(Figure 2A)**. TNFAIP2 and HCK were common DIU among anti-dsDNA^+^ and anti-dsDNA^+^ENA^+^ subsets, whereas TIMM10 was common in anti-ENA^+^ and anti-dsDNA^+^ENA^+^ subsets. FAM96A was common in all the SLE subsets **(Figure 2A)**. All the observed DIUs were the statistically significant (FDR p-value <0.05) **(Supplementary table 2)**. Interestingly, we observed that some of the DIU also demonstrates isoform switching. They were GRAP2 in anti-dsDNA^+^ patients **(Figure 2B)**; IGHM, OSCAR, ADORA2A, NACA and BRI3 in anti-ENA^+^ patients **(Figure 2C)** and AHCYL2 in anti-dsDNA^+^ENA^+^ patients **(Figure 2D)**.

**Figure 2.**
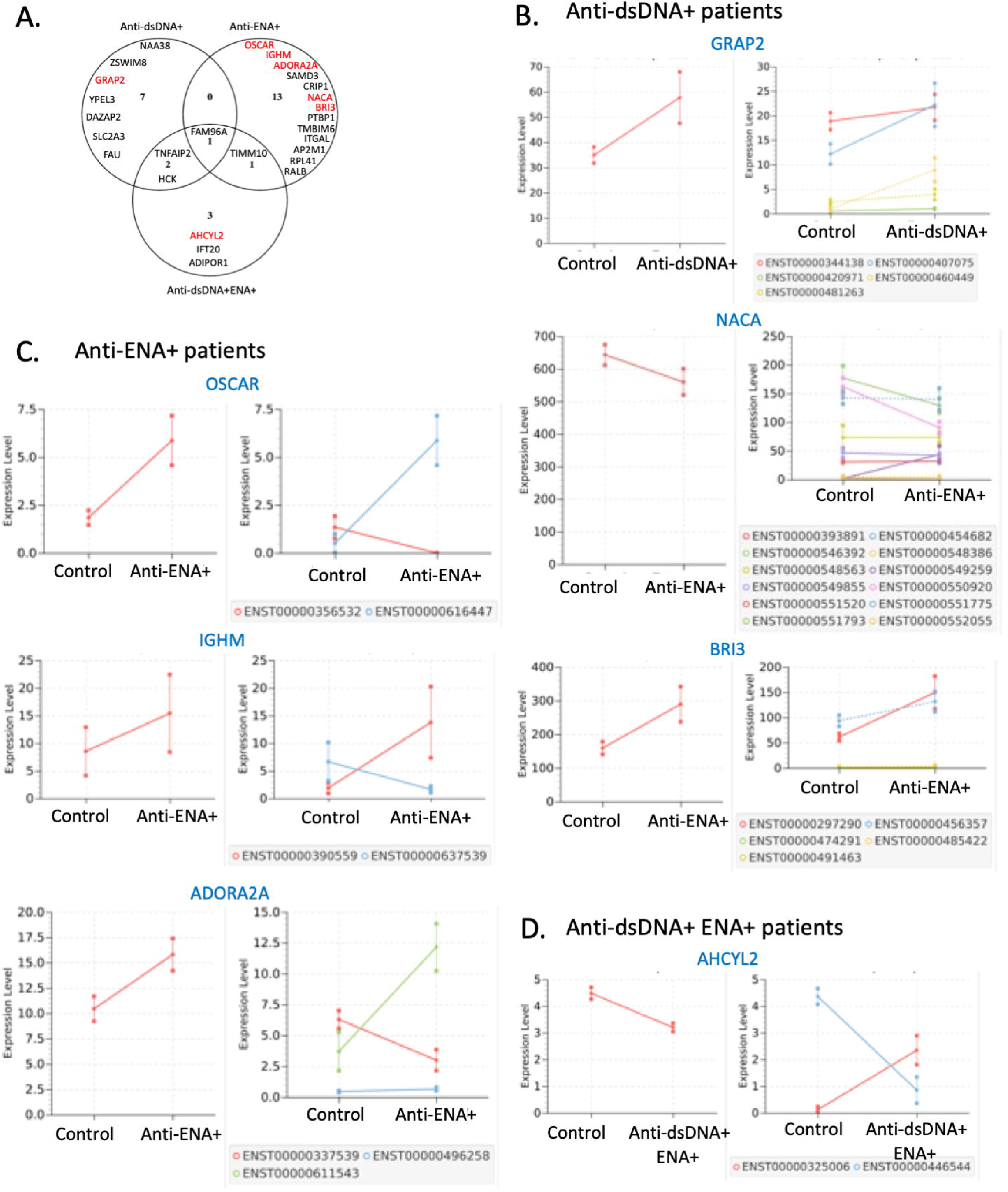
Distinct DIU genes in SLE patients’ subsets and comparison of their genes and isoform expression with controls. **A.** Venn diagram represents the distribution of unique as well as common DIU genes in distinct SLE patients’ subsets. The genes are arranged in the order of highest to lowest total usage change from top to bottom for unique DIU’s in each SLE subset. The “Red” colored gene are the gene which also shows isoform switching events **B.** GRAP2 in anti-dsDNA^+^ patients **C.** OSCAR, IGHM, ADORA2A, NACA and BRI3 in anti-ENA^+^ patients **D.** AHCYL2 in anti-dsDNA^+^ENA^+^ patients. *Each set of graph represents gene expression on the left which is combined expression of all isoforms demonstrating its variation in control and specific SLE patients’ subsets whereas, the graph on the right represents change in the isoform expression pattern of repective gene in control and SLE patients’ subsets.

In anti-dsDNA^+^ subgroup, only the isoforms of GRAP2 gene represent DIU as well as MIS. The major change in GRAP2 was the expansion of a retained-intron isoform ENST00000460449 in anti-dsDNA^+^ subgroup, it rises from about 3% to 15%. Another non-coding isoform ENST00000481263 along with ENST00000460449 isoform go from about 10% to 22% of the gene’s expression **(Figure 2B, Supplementary table 3)** indicating, it could be the main driver of the DIU signal. ENST00000344138 and ENST00000407075 were the two main protein-coding isoforms that encodes identical protein that give rise to O75791-1 (330aa). The expression of these protein-coding transcripts elevates about 40% (31.6-44.9), but its overall proportion falls from 90% to 78% **(Figure 2B, Supplementary figure 3A, Supplementary table 3)**. However, the transcript ENST00000420971 encoding a shorter protein B1AH86 (97aa) shows comparable change in control and anti-dsDNA^+^ subgroup **(Supplementary figure 3A)**. These isoform changes represent that the expression of GRAP2 was not suppressed while, the expression of isoforms was redistributed toward a non-productive isoform alongside higher overall expression.

Majority of the genes in anti-ENA^+^ SLE subgroups demonstrates DIU as well as MIS events. They were OSCAR, IGHM, ADORA2A, NACA and BRI3. OSCAR shows full reversal between the two protein-coding isoforms. The proportion of ENST00000356532 isoform of OSCAR changes from about 72% to 0%, while isoform ENST00000616447 goes from about 28% to 100% in anti-ENA^+^ subsets **(Figure 2C, Supplementary figure 3B, Supplementary table 3)**. The canonical isoform of OSCAR ENST00000284648 encoding protein Q81YS5 of 282aa was not detected in our cohort. Additionally, reversal of the dominant transcript was noted in IGHM gene. The transcript ENST00000637539 encoding protein P01871-2 of 474aa, express dominantly in controls (about 78%), whereas the transcript ENST00000390559 that encodes 453aa P01781-1 protein was profoundly expressed in anti-ENA^+^ subgroups with proportion of 89% **(Figure 2C, Supplementary figure 3B, Supplementary table 3)**. The proportion of canonical transcript ENST00000337539 of ADORA2A gene was found to be 60% and expressed dominantly in controls. In anti-ENA^+^ subgroup, it was replaced by ENST00000611543 at about 77%. The canonical transcript level diminished by 2-fold in anti-ENA^+^ subgroup, whereas the later one goes up about 3.3-fold, which was a reciprocal exchange, not a one-sided gain. Despite the isoform switch, they do not result in any protein changes as both encodes P29274 protein with 412aa. Another transcript, ENST00000496258 encoding protein S4R3A7 (83aa) was observed as low abundance but no change was noted **(Figure 2C, Supplementary figure 3B, Supplementary table 3)**. We observed twelve isoforms of NACA in anti-ENA^+^ subsets. The proportion of NACA gene transcript ENST00000549259 increases from 0.2% in control to about 8% anti-ENA^+^ subsets, on the other hand it was balanced mainly by decrease in the proportion of transcripts, ENST00000550920 and ENST00000546392 in same group. The former transcript encodes F8W0W4 (198aa protein), whereas the latter two encodes protein H0YHX9 (235aa) and Q13765-1 (215aa), respectively. Overall, the switch was noted in the protein-coding pool, but not in the non-coding genes. There was marginal variation in the processed transcript ENST00000551775 in anti-ENA^+^ subsets, from 25.0% in control to 23.1% in anti-ENA^+^ subsets **(Figure 2C, Supplementary figure 3B, Supplementary table 3)**. In controls, the major BRI3 transcript was the non-coding processed transcript ENST00000456357 about 59%, while in anti-ENA^+^ subgroup, the protein-coding canonical transcript ENST00000297290 encoding O95415-1 125aa (about 52%) representing dominant transcript reversal. The transcript ENST00000491463 encodes truncated protein 47 aa, as this is nonsense mediated decay transcript. Even though, it was elevated in anti-ENA^+^ subsets by log2FC 1.5, but it contributes to only 1% of the gene in both the groups **(Figure 2C, Supplementary figure 3B, Supplementary table 3)**.

In anti-dsDNA^+^ENA^+^ subgroup, AHCYL2 was the only gene that represent DIU and MIS. The dominant transcript of AHCYL2 reverses in anti-dsDNA^+^ENA^+^ subgroup. The transcript ENST00000446544 which encodes 610aa protein (Q96HN2-2) falls from about 97% to about 27% of the gene, whereas the ENST00000325006 transcript encodes 611aa protein (Q96HN2-1) which increases from about 3% to about 73% **(Figure 2D, Supplementary figure 3C, Supplementary table 3).** The transcript encoding 611aa was the canonical transcript which was nearly absent in controls. So, the switch was toward the annotated canonical form, not away from it which likely doesn’t affect the function. Overall, anti-dsDNA^+^ and anti-ENA^+^ patients showed predominant DIU however, isoform switching within the DIU genes was restricted to anti-ENA^+^ patient subset only. The remaining DIU events may reflect proportional transcript changes without a reversal in isoform dominance, suggesting regulatory modulation rather than classical isoform switching, with potential functional consequences.

### Functional consequences of identified transcriptional changes

To determine the functional relevance of the DIU changes with respect to each SLE patients’ subsets we used the combination of TUC percentage and the isoform fold change. Furthermore, we categorized these DIU gene with respect to changes from canonical isoform to protein coding or non-coding isoform in different SLE patients’ subsets **(Figure 3)**. It is noteworthy that, in all SLE subsets, most DIU genes give rise to altered protein-coding isoforms, resulting in either unchanged, elongated, or truncated protein sequences. These include DAZAP2, FAU, HCK, NAA38, TNFAIP2 and YPEL3 in anti-dsDNA^+^ subsets; ADORA2A, AP2M1, BRI3, CRIP1, IGHM, NACA, OSCAR, PTBP1, TIMM10 and TMBIM6 in anti-ENA^+^ subsets; and ADIPOR1, AHCYL2, HCK, IFT20 and TIMM10 in anti-dsDNA^+^ENA^+^ subsets **(Figure 3 and Supplementary table 3)**. Among these, isoform alterations in the protein-coding transcripts of ADORA2A, AHCYL2, AP2M1, CRIP1, IFT20, TIMM10 and TMBIM6 observed in specific SLE patient subsets, produce identical protein-coding isoforms, thereby likely preserving the associated biological functions without significant impact. Additionally, we observed that in distinct SLE subgroups, the canonical isoforms of FAM96A, GRAP2, SLC2A3, ZSWIM8, ITGAL, RPL41, RALB, SAMD3, TMBIM6, IFT20 and TNFAIP2 shifted to isoforms characterized by biotypes such as processed transcripts and/or retained introns **(Figure 3 and Supplementary table 3)**. This pattern suggests that post-transcriptional regulatory mechanisms may alter the functional properties of these genes. Notably, TMBIM6 in the anti-ENA^+^ subgroup and IFT20 in the anti-dsDNA^+^ENA^+^ subgroup exhibit isoform changes affecting both protein-coding and non-coding transcripts, highlighting complex regulation at the isoform level in SLE pathogenesis. **(Figure 3 and Supplementary table 3)**.

**Figure 3.**
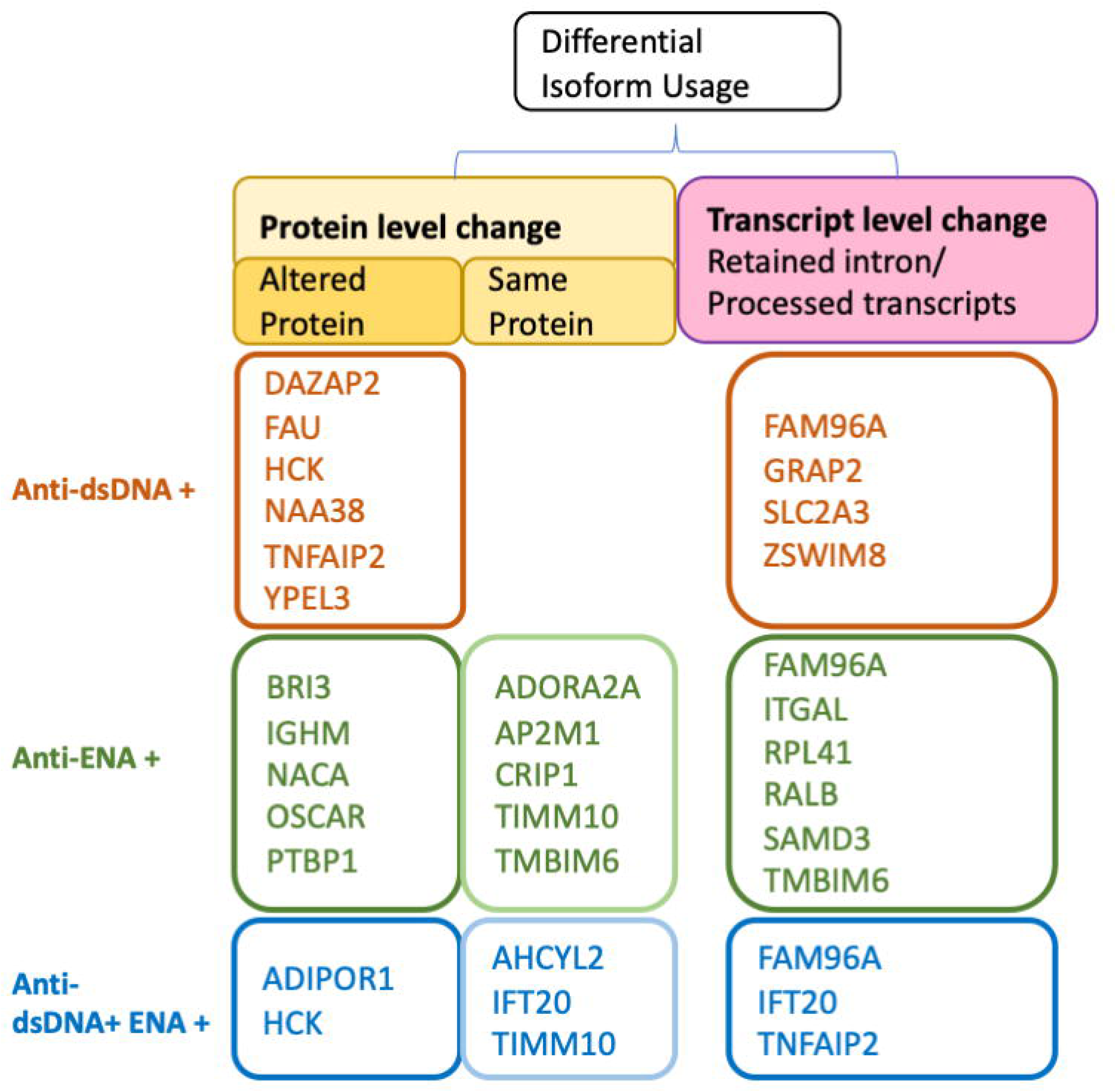
Illustration of DIU gene in all the SLE patients’ subsets. The dominant isoform (canonical isoform) result in either changes in the protein coding isoform (protein level change) or non-coding isoform (transcript level change).

To further explore the impact of alternative splicing in SLE, we quantified TUC across DIU genes in patient subsets. The extent of TUC among DIU genes varied considerably across SLE subgroups, ranging from 2% to 30% in the anti-dsDNA+ group, 4% to 72% in the anti-ENA^+^ group, and 5% to 70% in the anti-dsDNA^+^ENA^+^ group **(Supplementary table 2)**. This variability suggests differential isoform remodelling across SLE phenotypes. Given the relatively modest TUC values in the anti-dsDNA^+^ subgroup, a threshold of 20–30% was employed to determine biological relevance. In contrast, for the anti-ENA^+^ and anti-dsDNA^+^ENA^+^ subgroups, more stringent cut-offs >50% were applied to capture genes with potentially significant isoform shifts. Based on these criteria, we conducted focused analyses on NAA38 (TUC 30%) in the anti-dsDNA^+^ subgroup; OSCAR (73%) and IGHM (67%), in the anti-ENA^+^ subgroup; and AHCYL2 (71%) in the anti-dsDNA^+^ENA^+^ subgroup **(Supplementary table 2)**. ***(i) DIU in anti-dsDNA^+^ subgroup:*** Among the six unique DIU genes in anti-dsDNA^+^ patients, the NAA38 gene shows the maximum usage change of 30%. We observed six transcripts of NAA38, one processed transcript and five of those were protein-coding including the canonical isoform. The canonical isoform of NAA38, ENST00000575771 shows only slight downregulation in anti-dsDNA^+^ subgroup (logFC -0.15) compared to controls. This isoform encodes protein of 125aa residue, whereas two isoforms ENST00000575071 encoding 135aa residue protein and ENST00000575208 encoding 73aa residues protein (**Figure 4A**) were significantly overexpressed at logFC 1.96 (p <0.05) and 4.26 (p <0.05), respectively in anti-dsDNA^+^ subgroup (**Supplementary Table 3**). The later variant (ENST00000575208) represents use of an alternate promoter and therefore differs in the 5’ UTR and 5’ coding region compared to the other two variants and encode shorter isoform which has distinct N-terminus and lacks a predicted signal peptide compared to the canonical isoform and other isoform. Furthermore, to identify the functional impact of isoform switch of NAA38 we retrieved its interacting proteins from bioGRID (https://thebiogrid.org/) and performed the pathway analysis (**Figure 4B-C**). Interestingly, the top canonical pathway includes SLE, and top function is RNA degradation, mRNA splicing and spliceosome where primary interacting proteins are LSM2, LSM3, LSM4, LSM5, LSM6, LSM7 (**Figure 4C**). NAA38 was previously known as LSMD1. It forms NatC complex which localizes in cytoplasm and bind ribosomes. These proteins have a single Sm-like domain structure and associate with RNA and are often involved in RNA processing events [25, 26]. ***(ii) DIU in anti-ENA^+^ subgroup*:** In this subset, based on the TUC threshold OSCAR (72%) and IGHM (67%), was analyzed further. IGHM gene, encodes the heavy chain of IgM which has two isoform variants ENST00000637539 and ENST00000390559. The former one is the canonical isoform (encodes 474aa protein residue) is downregulated (logFC -2.03; p >0.05) whereas the later one is the switched isoform (encodes for 453aa residues) which was significantly elevated (logFC 2.88; p <0.05) in anti-ENA^+^ subsets. Among these isoforms the longer isoform encodes for membrane bound IgM (mIgM) whereas the shorter isoform encodes secretory IgM (sIgM), they differ only in the carboxy-terminal (C-terminal) region. The C-terminal of mIgM has hydrophobic transmembrane sequence and short cytoplasmic tail that is essential for anchoring on B cell surface, whereas this is totally absent in sIgM **(Figure 4D-E)**. OSCAR gene, encodes an osteoclast-associated receptor, which is a member of the leukocyte receptor complex protein family, a regulator of osteoclast differentiation which provides a co-stimulatory pathway. The canonical isoform ENSP00000365808 of OSCAR encodes a longer protein of 282aa residue. This isoform was not observed in anti-ENA^+^ subgroup instead we observed significant changes in the expression of other two isoforms ENST00000356532 (logFC -8.06; p<0.05) and ENST00000616447 (log FC 3.54; p<0.05) which could contribute to the changes in the isoform usage in anti-ENA^+^ subgroup. ENST00000356532 isoform encodes protein of 267aa. The crystal structure of the OSCAR revealed that it exists in both free and in complex with a consensus triple-helical peptide. The collagen-binding site is present in each immunoglobulin-like domain (D1 and D2) [27, 28]. The canonical isoform encodes secretory OSCAR (OSCAR-S), whereas the later isoform is the membrane bound OSCAR (OSCAR-M), it has a type I transmembrane protein of 263 amino acids (245 after signal peptide removal), with two Ig-like domains D1 and D2 in its N-terminal extracellular region, a predicted transmembrane region, and a short C-terminal cytoplasmic tail **(Figure 4F-G)**. ***(iii) DIU in anti-dsDNA^+^ENA^+^ subgroup:*** In this subset, only AHCYL2 shows the maximum TUC of 71%. AHCYL2 gene encodes AHCY-like family of protein which has high sequence homology with S-adenosyl homocysteine hydrolase (AHCY). Due to 97% homology with AHCY domain it is suggested to be involved in methionine cycle which involves the conversion of S-adenosyl-L-homocysteine to L-homocysteine and adenosine. Among the two isoforms observed in our study, the canonical isoform of AHCYL2, ENST00000325006 is 4-fold upregulated (p<0.05) in anti-dsDNA^+^ENA^+^ group whereas the ENST00000446544 isoform is 2-fold downregulated (p<0.05). The former one canonical isoform encodes 611aa protein whereas the later one encode a protein of 610aa residue. These two isoform encodes protein that differs only in one amino acid at 122aa position at the N-terminal which was neither highly conserved nor involved in catalytic functions **(Supplementary figure 4)**.

**Figure 4.**
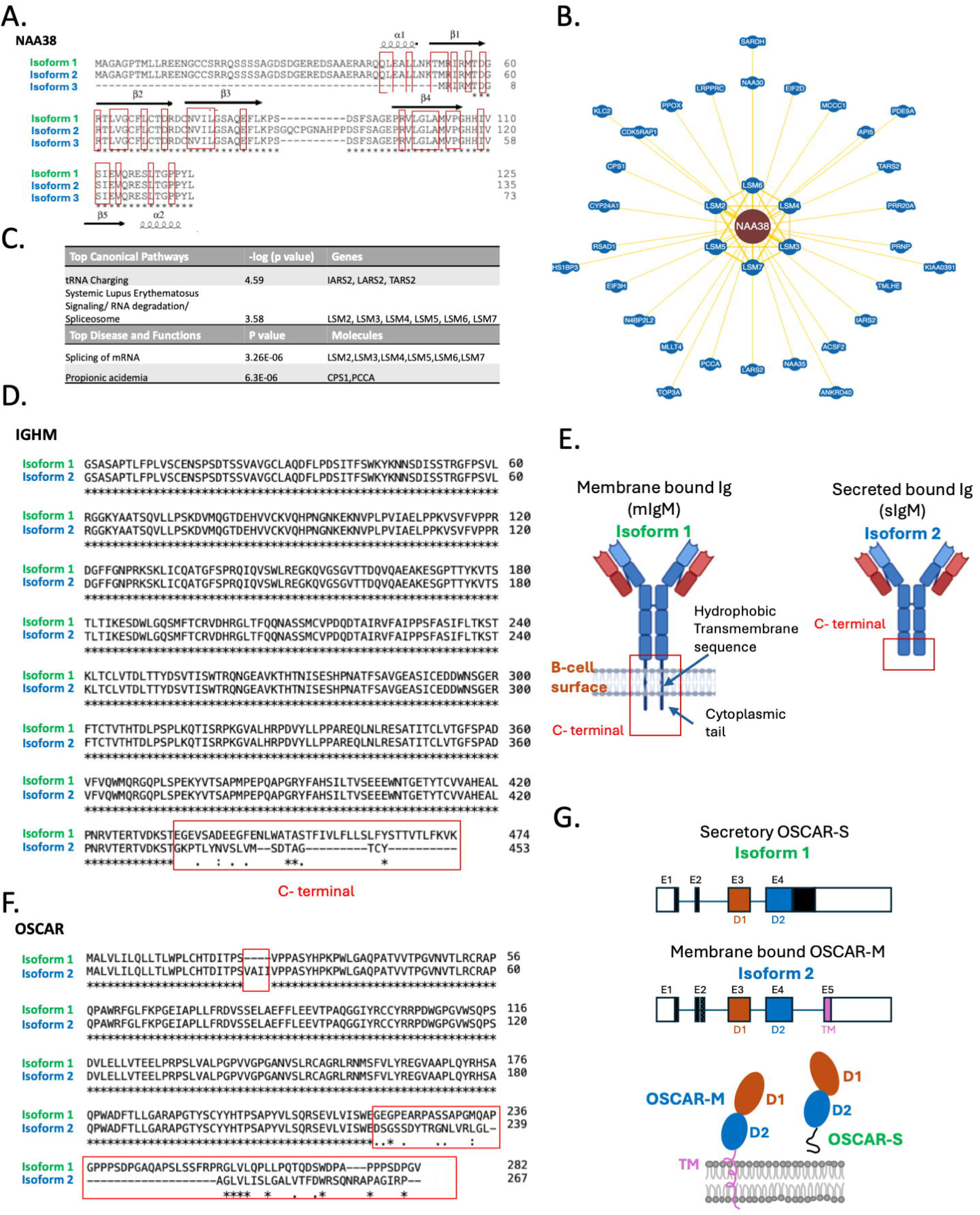
Isoform description and sequence alignment of NAA38, IGHM and OSCAR proteins. **A.** NAA38 canonical (green) and non-canonical (blue) isoforms **B.** Network of proteins interacting with NAA38 **C.** Pathway analysis of the NAA38 interacting protein **D.** IGHM protein sequence derived from canonical (green) and non-canonical (blue) isoforms E. Structural difference in the mIgM and sIgM **F.** OSCAR protein sequence derived from canonical (green) and non-canonical (blue) isoforms **G.** Structural difference in the secretory and membrane bound OSCAR.

### Isoform level functional diversity analysis and functional feature inclusion in DIU genes in SLE patients’ subsets

To assess the isoform-level or protein-level structural variation of a specific DIU gene within the SLE patients’ subsets, we performed functional diversity analysis. We noted the transcript level variation in approximately 90-100% of the annotated DIU genes across following categories: 3′UTR length, 5′UTR length, CDS length, PAS, uORF, miRNA-binding sites, repeat elements, in all SLE patients’ subgroups, except for OSCAR and AHCYL2 gene transcript, in anti-ENA^+^ and anti-dsDNA^+^ENA^+^ patients’ subsets, respectively **(Figure 5A)**. In contrast, variation in protein-level features was low overall, but the degree of variations differs in each SLE patients’ subset. We observed signal peptide variation in anti-ENA^+^ patients only at 33%. Notable variation in PTM sites was found in anti-dsDNA^+^ (75%), anti-ENA^+^ (82%) and anti-dsDNA^+^ENA^+^ (33%). OSCAR and AHCYL2, both did not show any diversity even at protein-level **(Figure 5A)**. The transcripts that demonstrate the variation in the transcript and protein were queried for the differential feature inclusion.

**Figure 5.**
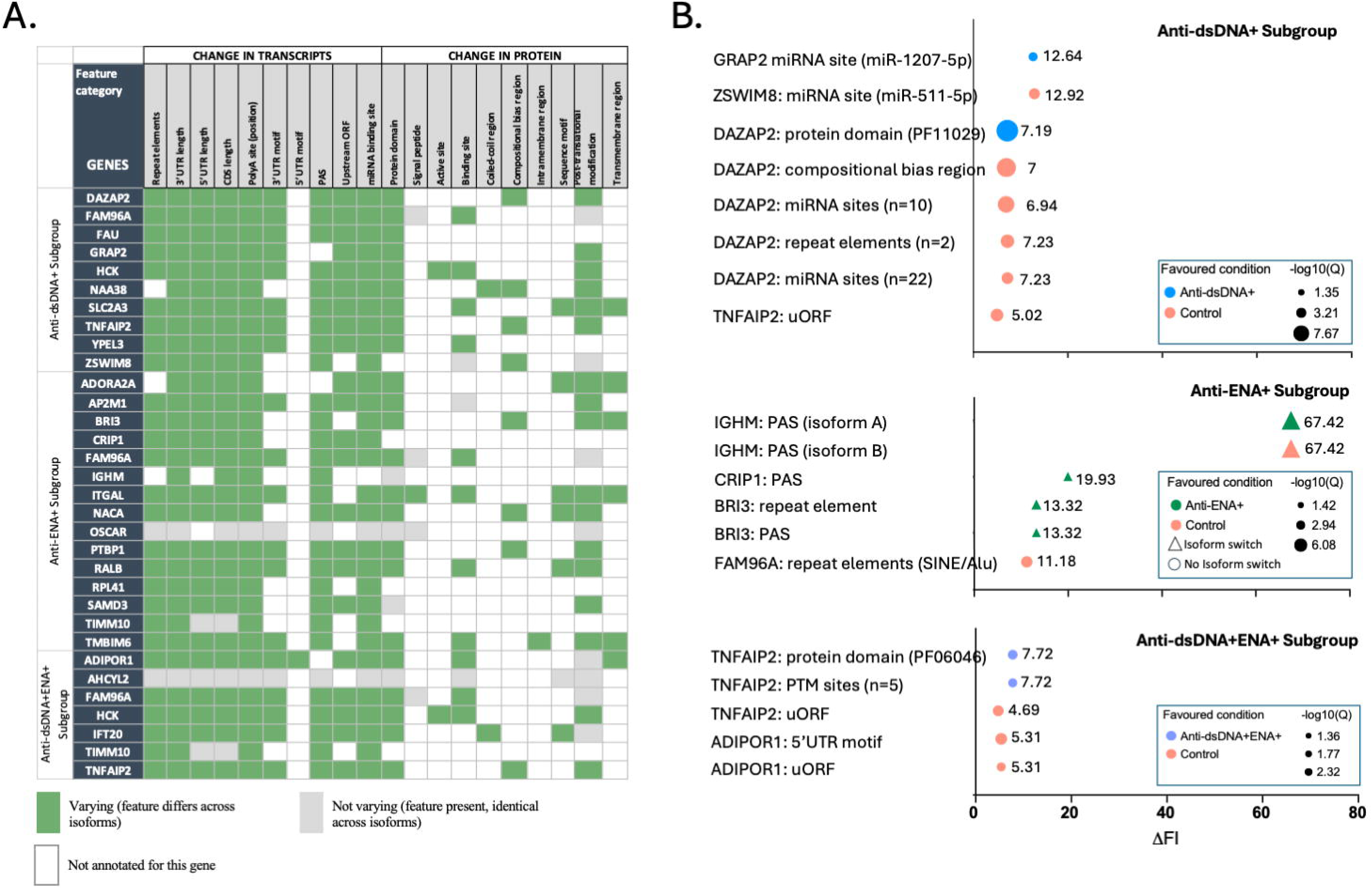
Functional diversity analysis (FDA) of transcript- and protein-level features and differential feature inclusion across all the SLE patients’ subset A. For each gene, cells indicate whether an annotated feature category differs *(“varying” as green)* or is identical *(“not varying” as grey)* or not annotated *(“blank” as white)* across the gene’s transcript/protein isoforms, independent of expression or disease-group comparison. Genes shown were prioritized for downstream differential feature inclusion (DFI) testing. B. The dot plot demonstrates DFI of the DIU gene in anti-dsDNA^+^ (top) anti-ENA^+^ (middle) and anti-dsDNA^+^ENA^+^ (bottom); ΔFI (effect size on the x-axis; Color= favoured condition (blue= anti-dsDNA^+^, green=anti-ENA^+^, purple=anti-dsDNA^+^ENA^+^ and red=controls), Marker shape = triangle-major isoform switch, circle-no isoform switch; Marker size = statistical significance (larger –smaller Q value, in the figure represented as –log10(Q)).

Furthermore, we assessed the functional feature inclusion in isoforms of observed DIU genes and investigated if these events were favored in specific SLE patients’ subsets or controls. We observed significant differential feature inclusion in GRAP2 and DAZAP2 in anti-dsDNA^+^ subsets; IGHM, BRI3 and CRIP1 in anti-ENA^+^ subset; and TNFAIP2 in anti-dsDNA^+^ENA^+^ subset compared to healthy individuals **(Figure 5B)**. ENST00000460449 of GRAP2 which is a non-protein coding isoform, shows inclusion of miR-1207-5p target site which suggest that the gene expression of GRAP2 is regulated post-transcriptionally by this miRNA. DAZAP2 transcript, ENST00000551313 shows a significant (Q = 2.14x10^-8^) protein (PF11029) domain inclusion favoring anti-dsDNA^+^ subsets, while the transcript ENST00000412716 with proline-rich compositional-bias inclusion and a longer transcript harboring an extensive 3′ region encompassing 32 annotated miRNA-binding sites (spanning members of the miR-27, miR-378, miR-520/525 and other families) and two simple-repeat elements, were preferentially retained in controls **(Figure 5B and Supplementary table 4)**. The protein encoded by ENST00000551313 transcript is F8VUW5 (108aa) whereas the canonical transcript ENST00000412716 encodes Q15038-1 protein (168aa). The shorter protein lacks key elements SH2 domain **(Supplementary figure 5A, Supplementary table 5)**. Loss of a miR-511-5p binding site in ZSWIM8 transcripts ENST00000451629, and loss of an uORF in TNFAIP2 isoform ENST00000558056 in anti-dsDNA^+^ subsets favours controls. A total of eight transcripts of TNFAIP2 was observed in anti-dsDNA^+^ **(Supplementary table 3)**. ENST00000561217 non-coding processed transcript stays dominant at about 35% in both controls and anti-dsDNA^+^ subsets. The short uORF-bearing ENST00000558056 transcript encoding 39aa protein was depleted by about 5 percentage in anti-dsDNA^+^ patients, thus favouring controls. Overall, gene expression of TNFAIP2 was observed to be lower by 37% in ant-dsDNA^+^ subgroups. ZSWIM8 in the anti-dsDNA^+^ subset, showed DIU without major isoform switching, driven by non-coding transcripts, most notably near-complete loss of the processed transcript ENST00000451629 from 13.2% to 0.3% **(Supplementary table 3)** whereas, the fraction of protein-coding transcripts ENST00000604729 and ENST00000605216 remains dominant at 38% and at comparable level in both controls and anti-dsDNA^+^ subsets. This indicates that the loss of the miR-511-5p binding site favours controls is the feature-level consequence of losing that transcript, not a separate regulatory change.

Interestingly, we observed most striking signal in IGHM in anti-ENA^+^ subgroup, the poly adenylation site (PAS) inclusion in IGHM transcript ENST00000390559 (ΔFI = 67, Q=8x10^-7^), whereas the alternate transcript ENST00000637539 favoring controls (ΔFI = 67, Q=8x10^-7^). CRIP1 transcript ENST00000392531 and BRI3 transcript ENST00000297290 shows PAS driven isoform switches, both favoring in anti-ENA^+^ subgroup. Parallely, ENST00000297290 isoform of BRI3 demonstrates repeat-element-driven isoform switches. However, the protein coding isoforms in CRIP1 translates to same protein P50238 (77aa). FAM96A transcript shows inclusion of SINE/Aly repeat element favouring control but not a major isoform switch **(Figure 5B and Supplementary table 4)**.

Nine TNFAIP2 transcripts were expressed in anti-dsDNA^+^ENA^+^ subgroup, ENST00000333007 transcript represents four distinct post-translational modification such as acetylation, phosphorylation of serine, threonine and tyrosine, whereas another transcript, ENST00000560670 shows inclusion of protein domain (PF06046). Compared to anti-dsDNA^+^ENA^+^, we found loss of an upstream open reading frame (uORF) in ENST00000558056 isoform in controls **(Supplementary table 4)**. TNFAIP2 shows subtle isoform usage change but no the isoform switching events **(Supplementary table 2)**. As PTM sites in ENST00000333007 favour anti-dsDNA^+^ENA^+^ subsets, combined proportion of transcripts ENST00000333007 and ENST00000560869 that encodes same protein Q03169 (654aa) was 85% of protein-coding expression in controls, while 90% in anti-dsDNA^+^ENA^+^ subsets, mainly because the expression of transcript ENST00000558056 (encoding 39aa protein) reduced from 24.1 to 9.6 and ENST00000560869 rises from 60.6 to 90.4. The proportion of transcript with PF06046 domain favouring anti-dsDNA^+^ENA^+^ subsets was twice as much as that of transcript in controls. The level of ENST00000558056 transcripts that encodes 39aa protein drops in anti-dsDNA^+^ENA^+^ subsets from 7.2% to 2.5%, they do not contain the uORF domain in these subsets **(Supplementary table 3)**. Overall, the total expression of TNFAIP2 gene was found to be higher in anti-dsDNA^+^ENA^+^ subsets by 14% which is reverse to that was observed in anti-dsDNA^+^ subsets. ADIPOR1 transcript ENST00000340990, demonstrates loss of uORF as well as 5’UTR motif in favouring controls **(Figure 5B and Supplementary table 4)**. ADIPOR1 also showed DIU without major isoform switching where the full-length transcript ENST00000340990 encoding 375aa protein remains dominant in both controls and anti-dsDNA^+^ENA^+^ subsets as 97% and 92%, respectively. A subtle increase in a truncated protein-coding transcript ENST00000426229 was found in anti-dsDNA^+^ENA^+^ subsets from 1.7% to 6%. The transcript ENST00000426229 encode protein of 205aa which has only two transmembrane domains compare to its canonical counterpart **(Supplementary figure 5B-C)**.

## Discussion

Gene expression profiling in SLE patients either as a “single group” or “distinct subsets based on autoantibody repertoire” helped in revealing the comprehensive biology of disease [1, 6, 7, 17, 18]. These studies have uncovered key pathways and gene expression signatures associated with disease activity and clinical heterogeneity. However, genes exhibiting isoform switching and differential isoform usage change, reflects a transcript level regulatory mechanism that remains underappreciated in SLE. To address this gap, the present study undertakes a comprehensive analysis of DIU genes, aiming to explore one of the most dynamic and functionally relevant aspects of transcriptome biology.

GO analysis of subset specific MIS genes highlighted enrichment of RNA-binding, protein binding, ubiquitin-like protein binding, and RNA polymerase II promoter binding pathways in either one or more SLE patient subsets. Specifically, association of anti-ENA^+^ autoantibodies with the spliceosome components is very well established [29, 30]. Recent studies have also shown that an alternative exon usage in TRIM21 generates unique Ro52 isoforms that are preferentially targeted by anti-Ro52 antibodies in SLE, illustrating how splicing variants can directly generate novel antigenic epitopes [31]. Evidence of anti-ENA autoantibodies (e.g. anti-RNP and anti-Sm) targeting core components of the spliceosome [30, 32], also points towards immune targeting of RNA-binding proteins involved in splicing which could contribute to more DIU or MIS events specifically in the anti-ENA^+^ patient subgroup. However, direct link of anti-dsDNA^+^ autoantibodies with protein binding or splicing events is not documented till date.

Differential isoform usage genes, though fewer in number, hold particular interest due to their potential to encode distinct protein isoforms with altered or novel functions. Genes with differential isoform usage demonstrating isoform switching in GRAP2 (anti-dsDNA^+^ subsets); IGHM, OSCAR, ADORA2A, NACA, BRI3 (anti-ENA^+^ subsets); and AHCYL2 (anti-dsDNA^+^ENA^+^ subsets) as well as those without isoform switching events were observed in each patient subset with a predominance in anti-ENA^+^ subsets. The variable TUC across patient subsets reveals differential extents of isoform remodeling, with more pronounced shifts in anti-ENA^+^ and anti-dsDNA^+^ENA^+^ subgroups. This quantitative insight guided our focused analysis on genes with biologically significant TUC values, providing candidate genes such as NAA38, OSCAR, IGHM, ADORA2A, and AHCYL2 for further functional investigation. Notably, many of the DIU genes identified in this study have not been previously implicated in SLE or other autoimmune conditions, indicating unexplored regulatory mechanisms. GRAP2 is an adaptor protein containing an N-terminal SH3 domain, a central SH2 domain, and a C-terminal SH3 domain. It acts as a molecular bridge for the T-cell receptor (TCR) signaling and co-stimulatory activation [33, 34]. GRAP2 isoforms in anti-dsDNA^+^ shows expansion of non-coding transcript and redistribution between two transcripts encoding the identical protein, indicating altered transcript composition rather than replacement of the protein product. Differential inclusion of a miRNA-binding site was favoured in anti-dsDNA^+^ subset of SLE patients. There are no direct evidence linking GRAP2 to SLE pathogenesis but its role in T cell signaling [33] and implication of aberrant T cell signaling in SLE [35] indicates that these events may influence the level or regulation of GRAP2 available to propagate TCR signalling. NAA38 did not show significant isoform switching but show the modest usage change in patients with anti-dsDNA^+^ autoantibody. NAA38 constitutes the small auxiliary subunit of the evolutionarily conserved NatC complex, along with the catalytic subunit NAA30, and the ribosomal anchor NAA35. The NatC complex mediates the N-terminal acetylation of proteins, a modification that can influence protein stability by protecting them from degradation. Varland et al., demonstrated that NAA38 is essential for optimal NatC activity [36]. This suggests that isoform usage change in NAA38 could result in the expression of a truncated or altered protein variants, potentially impacting downstream processes such as ubiquitination. While alterations in NAA30 have been implicated in several diseases [37], the role of NAA38 in SLE remains largely unexplored. Although, NAA38 is also known as LSMD1 and contains an Sm-like domain, a structural motif found in proteins associated with the ENA complex, but there is currently no direct evidence linking NAA38 to SLE.

In anti-ENA^+^ subgroup, both IGHM and OSCAR show MIS as well as DIU with higher total usage change suggesting reversal of the dominant isoforms compared to the controls. We observed the IGHM isoform that encodes sIgM protein was elevated in anti-ENA^+^ subsets. Studies from B cell IgM mRNA suggest that chains are identical except for their 41 and 20 residue C-terminal segments, respectively [38, 39]. Additionally, higher secretory IgM indicates stronger plasma cell signature [40]. In previous study, we reported plasma cell signature genes expressed in anti-dsDNA^+^ and anti-ENA^+^ SLE patient’ subsets [1]. In autoimmune diseases such as SLE, alterations in IgM isoform expression or function may impair clearance of self-antigens, thereby contributing to immune dysregulation and the generation of pathogenic autoantibodies [41–43]. Even though, the two isoforms of OSCAR represent MIS as well as DIU with highest TUC, but they encode the same membrane bound protein OSCAR-M. However, the canonical isoform of OSCAR gene that encodes secretory OSCAR-S protein which could not be detected. The wider expression of OSCAR by human myeloid immune cells suggested that OSCAR might play a role in innate immunity in addition to the reported role in osteoclastogenesis [44, 45]. However, no feature inclusion and functional diversity was observed in OSCAR transcript and protein, suggesting minimal impact in these SLE patients. Similarly, ADORA2A produce multiple transcripts that were switched in controls and anti-ENA^+^ patient subgroups, yet they both translates into the exact same protein. Interestingly, we observed redistribution among NACA protein-coding isoforms without a change in overall gene expression which indicates that isoform-level analysis detects regulation that gene-level expression testing would have missed. NACA gene encodes the alpha subunit of the nascent polypeptide-associated complex (NAC), a highly conserved cytosolic factor that operates primarily as a ribosome-bound molecular chaperone [46, 47]. The NACA was evident as an auto allergen that binds to IgE from atopic dermatitis patients’ sera and had been shown to induce the T cell proliferation and IFN-secreation [48]. Our observation of shift in the protein-coding isoform composition in anti-ENA^+^ subset suggests that this switch could determine which NACA epitopes were presented to the immune cells. The altered NACA isoform composition may relate to the switch IGHM isoform encoding sIgM, and isoform specific differences in NACA could modify the epitopes available for autoantibody recognition indicating its plausible implication in SLE. The proportion of protein-coding isoform as well as the total gene expression of BRI3 was observed to be elevated, the net effect was more profound towards BRI3 protein-coding transcript in anti-ENA^+^ subsets. Functionally, BRI3 participates in TNF-induced cell death and shown to be upregulated during TNF-mediated inflammation [49]. Thus, increased BRI3 protein-coding output could reflect its contribution towards altered TNF-related cell-death and inflammatory. To our knowledge, BRI3 have not been studied in SLE, however, dysregulated apoptosis and TNF-α are implicated in SLE [1, 18].

AHCYL2 in anti-dsDNA^+^ENA^+^ subgroup demonstrates isoform switch and maximum usage change. Both the transcripts differ by 30 bp, but protein changes by one aa residue only, so the remaining ∼27 bp lies outside the coding region, most likely in the UTRs or an alternative exon boundary indicating that no major changes in the protein isoform. Functionally, AHCYL2 is involved in one-carbon metabolism and amino acid pathways, particularly the conversion of S-adenosyl-L-homocysteine (SAH) to L-homocysteine and adenosine. SAH is a harmful by-product, its accumulation inhibits autophagy therefore, its hydrolyzed as soon as these are formed [50]. However, like OSCAR in anti-ENA^+^ subgroup, no feature inclusion and functional diversity was observed in AHCYL2 transcript and protein, suggesting minimal impact on the SLE biology. Taken together, these findings imply co-existence of different layers of regulatory mechanisms in each SLE patient subset. DIU without isoform switching likely reflects fine-tuning of transcript composition through splicing regulation or differential isoform stability, rather than functional replacement of one isoform by another unlike isoform switching [22, 51].

The diversity analysis and functional feature inclusion reveals inclusion of significant features in some of the DIU genes across different subsets of SLE patients, underscoring the complexity of gene regulation in autoimmune diseases. The data suggest each autoantibody subgroup carries a distinct isoform regulatory signature: subtle, non-switching regulatory changes in anti-dsDNA^+^ disease; a coordinated, switching event driving effects on immunoglobulin and eventually B-cells and 3′-UTR processing in anti-ENA^+^ subgroup of SLE patients; and a insertion of protein domain as well as PTM altering isoform change concentrated on TNFAIP2 in anti-dsDNA^+^ENA^+^ subset. Specifically, we observed post-transcriptional regulation of GRAP2 through the inclusion of miRNA binding sites, suggesting that its expression is fine-tuned by specific miRNAs in the anti-dsDNA^+^ subset. GRAP’s role in T cell receptor signaling [33, 34]. DAZAP2 is an RNA-binding/Wnt-signaling regulator, the inclusion of protein domains in DAZAP2 reflects functional alterations that may influence its RNA-binding activity, potentially impacting immune regulation in the same SLE subgroup. DAZAP2 is a ubiquitously expressed protein [52] which has a role in transcriptional regulation of TCF gene [53] as well as interacts with eukaryotic initiation factor 4G [54], ubiquitin protein ligase (E3) [55]. The shorter isoform shows a significant protein domain inclusion, whereas it has some missing key elements which could impact their functional properties as described above. The loss of a miR-511-5p binding site in ZSWIM8 (favoured in controls) corresponded to the marked reduction of the non-coding transcript ENST00000451629 in anti-dsDNA^+^ patients, indicating that the DFI and DIU results reflect a single transcript level event that does not affect protein-coding output. Strikingly, IGHM demonstrates major isoform switching alternative 3′ processing of IGHM controls the switch between the secreted vs membrane-bound forms of IgM, so this finding may reflect altered B-cell antibody secretion dynamics in anti-ENA^+^ subsets [41–43]. CRIP1 and BRI3 show parallel PAS and repeat element-driven isoform switches, both favoring the anti-ENA^+^ subgroup, suggesting a broader pattern of altered 3′-UTR processing based isoform usage in this subgroup rather than an isolated event. Additionally, PAS inclusion in IGHM, CRIP1, and BRI3 in the anti-ENA^+^ subset suggests changes in mRNA stability and translation efficiency, which may contribute to altered B-cell function and immune response. Despite differential feature inclusion in CRIP1 isoform, the changes were not translated at the protein level suggesting the minimal effect in impacting the disease pathology. Similarly, in ADIPOR1 in anti-dsDNA^+^ENA^+^ subsets, the DFI of uORF and the 5′UTR motif were on the dominant transcript favouring controls alongside a small increase in a truncated protein-coding transcript, indicating subtle transcript-level modulation. The observed minimal change with 5% decrease in anti-dsDNA^+^ENA^+^ subsets was unlikely to cause any de-repression events for this gene. TNFAIP2 is the TNF-α inducible protein 2, it was primarily identified as an inducible gene in response to TNF-α, also called as B94 or M-Sec [56]. TNFAIP2 is best characterised as a driver of tunneling nanotube formation in macrophages, which serve as a communication between myeloid cells, immune cells under pro-inflammatory activation [57, 58]. Both anti-dsDNA^+^ and anti-dsDNA^+^ENA^+^ subsets showed depletion of the short uORF-containing transcript ENST00000558056 (∼7% to ∼2–3%). In the anti-dsDNA^+^ENA^+^ subset, TNFAIP2 showed DIU without major isoform switching, yet the accompanying DFI results indicate a coordinated shift in regulatory features, like carrying phosphorylation and acetylation sites (ENST00000333007) and a PF06046 domain-containing isoform (ENST00000560670) were favoured in this subset, whereas a uORF-containing isoform was favoured in controls. The redistribution of TNFAIP2 transcripts we observed in the anti-dsDNA^+^ENA^+^ subset, toward full-length coding isoforms carrying phosphorylation and acetylation sites and away from a uORF-containing isoform, could therefore increase TNFAIP2 protein output and its responsiveness to post-translational signalling in myeloid cells. Although, total gene expression was reduced in anti-dsDNA^+^ (∼37%) but modestly increased in anti-dsDNA^+^ENA^+^ (∼14%), indicating subgroup-specific regulation of the full-length transcripts. Notably, this isoform-level regulation in anti-dsDNA^+^ENA^+^ subsets occurs without any change in the dominant transcript and with only a modest change in total gene expression, it would not be detectable in conventional gene-level analyses. This is consistent with the broader hypothesis that different SLE autoantibody profiles reflect distinct post-transcriptional regulatory programs rather than a single shared mechanism. OSCAR and AHCYL2 did not exhibit any kind of functional diversification (through splicing changes, domain modifications, or novel isoform generation) which suggests that the gene’s expression, splicing, and protein features remain consistent between SLE patients’ subsets and controls. As a result, it may not be directly involved in the pathophysiology of SLE. Taken together, understanding how functional diversity varies in diseased condition can help in identifying therapeutic targets, particularly in autoimmune diseases, cancer, and chronic inflammatory conditions.

Together, our findings indicate that autoantibody defined SLE subgroups differ in isoform usage in ways that were largely neglected at the gene level analysis. Many isoform shifts occurred between transcripts encoding identical or near-identical proteins, and the associated DFI features (PAS, uORFs and miRNA binding sites) point to altered post-transcriptional control as the main functional consequence. The affected genes converge on lymphocyte signalling and antibody secretion (GRAP2, IGHM), TNF-related pathways (BRI3, TNFAIP2) and RNA regulation (ZSWIM8, DAZAP2), with the strongest switching in the anti-ENA^+^ subgroup. Whether these changes reflect altered cell composition or regulation within cells remains to be determined; future work using long-read and cell-type-resolved sequencing, protein-level validation and independent clinical cohorts will be needed to establish their functional and diagnostic relevance.

## Authors’ contribution

RR and GR conceived and designed the study. RR performed the experiments, and RR, AD, SKC, VVS, DV and GR analyzed and interpreted the data. MR provided the SLE patient samples. RR and GR wrote the paper. GR critically reviewed and revised the manuscript, provided overall guidance for the project and approved the final manuscript as submitted. All authors read and approved the final manuscript.

## Aknowledgements

This study was supported by a grant (BT/PR4619/MED/30/834/2012) from the Department of Biotechnology, New Delhi, India to GR. RR was supported by senior research fellowship from Council of Scientific & Industrial Research (CSIR), New Delhi, India. The funding agency had no role in study design, data collection, analysis and decision to publish the paper.

## Supplementary Data

**Supplementary Figure 1. Illustration of isoform switch and isoform usage change in diseased condition A.** Representation of multiple isoforms of a gene wherein isoform 1 is the most dominant isoform also known as canonical transcript **B.** A graph depicts an example of isoform switch in healthy and diseased group, isoform 1 is dominantly expressed in healthy whereas isoform 2 is predominantly present in diseased condition. The magnitude of the isoform usage change from isoform 1 to isoform 2 in diseased condition is defined as TUC.

**Supplementary Figure 2. Distribution and Gene ontologies for MIS in each SLE patient subsets A.** The number of gene that undergoes post-transcriptional modification (DIU and IS) and differential gene (DE) regulation in different subsets of SLE patients **B.** Venn diagram represents the distribution of unique as well as common genes with MIS among distinct subsets of SLE patients **C**. Gene ontologies term enriched for unique MIS in each SLE subsets.

**Supplementary Figure 3. Expression of Protein coding isoforms A.** Anti-dsDNA^+^ subsets-GRAP2 **B.** Anti-ENA^+^ subsets-OSCAR, NACA, IGHM, BRI3, ADORA2A **C.** Anti-dsDNA^+^ENA^+^ subsets-AHCYL2.

**Supplementary Figure 4. Sequence alignment of AHCYL2 protein A.** AHCYL2 protein sequence derived from canonical and non-canonical isoforms **B.** Similarity in the AHCYL2 domain of two isoforms.

**Supplementary Figure 5. Sequence** alignment of DAZAP2 and ADIPOR1 protein **A.** DAZAP2 protein sequence derived from canonical and non-canonical isoforms **B.** ADIPOR1 protein sequence derived from canonical and non-canonical isoforms **C.** Structural difference in the transmembrane domain of ADIPOR1in canonical and non-canonical isoform. *TM-Transmembrane, E-Extracellular, C-Cytoplamsic domain

**Supplementary Table 1.** List of major isoform switch genes in each SLE patients’ subset

**Supplementary Table 2.** Differential isoform usage analysis across SLE patients’ subsets

**Supplementary Table 3.** Isoforms of DIU genes

**Supplementary Table 4.** Transcript- and protein-level features showing significant differential feature inclusion in DIU gene across SLE patients’ subsets

**Supplementary Table 5.** Known interaction partners of DAZAP2 and presence of the corresponding binding regions in the canonical and altered isoform

