## Supplementary material for "Preferential Isoform Usage and Isoform Switching in Systemic Lupus Erythematosus Patients with Distinct Autoantibody Profile": Supplentary table 2

**Table 2. Differential isoform usage analysis across SLE patients’ subsets**

| **Gene** | **Gene description** | **Q value** | **Major isoform switching** | **Total usage change (%)** | **Isoforms** | **Control mean expression** | **Disease-group mean expression** |
| --- | --- | --- | --- | --- | --- | --- | --- |
| **Control vs. Anti-dsDNA+ SLE subset** | | | | | | | |
| ***DAZAP2*** | DAZ associated protein 2 | *< 0.0001* | No | 7.48 | 8 | 372.00 | 335.76 |
| ***TNFAIP2*** | TNF alpha induced protein 2 | 0.0012 | No | 9.58 | 8 | 329.02 | 208.80 |
| ***SLC2A3*** | solute carrier family 2 member 3 | 0.0014 | No | 5.55 | 8 | 130.22 | 219.25 |
| ***ZSWIM8*** | zinc finger SWIM-type containing 8 | 0.0027 | No | 18.72 | 12 | 37.01 | 32.67 |
| ***FAU*** | FAU, ubiquitin like and ribosomal protein S30 fusion | 0.0044 | No | 2.09 | 6 | 708.21 | 752.19 |
| ***YPEL3*** | yippee like 3 | 0.0110 | No | 11.11 | 6 | 276.52 | 239.66 |
| ***FAM96A*** | family with sequence similarity 96 member A | 0.0195 | No | 12.42 | 5 | 36.02 | 34.25 |
| ***GRAP2*** | GRB2-related adaptor protein 2 | 0.0257 | Yes | 16.47 | 5 | 34.94 | 57.74 |
| ***HCK*** | HCK proto-oncogene, Src family tyrosine kinase | 0.0280 | No | 5.93 | 5 | 224.11 | 163.80 |
| ***NAA38*** | N(alpha)-acetyltransferase 38, NatC auxiliary subunit | 0.0412 | No | 30.18 | 6 | 19.39 | 24.12 |
| **Control vs. Anti-ENA+ SLE subset** | | | | | | | |
| ***IGHM*** | immunoglobulin heavy constant mu | 1.59 × 10^-7^ | Yes | 67.42 | 2 | 8.53 | 15.42 |
| ***NACA*** | nascent polypeptide-associated complex alpha subunit | 2.24 × 10^-4^ | Yes | 13.78 | 12 | 643.10 | 560.09 |
| ***OSCAR*** | osteoclast associated, immunoglobulin-like receptor | 0.0118 | Yes | 72.59 | 2 | 1.84 | 5.87 |
| ***BRI3*** | brain protein I3 | 0.0141 | Yes | 13.44 | 5 | 159.20 | 289.36 |
| ***ADORA2A*** | adenosine A2a receptor | 0.0197 | Yes | 41.42 | 3 | 10.44 | 15.80 |
| ***FAM96A*** | family with sequence similarity 96 member A | 0.0027 | No | 14.10 | 5 | 37.03 | 37.23 |
| ***TIMM10*** | translocase of inner mitochondrial membrane 10 | 0.0027 | No | 23.75 | 3 | 18.84 | 76.57 |
| ***SAMD3*** | sterile alpha motif domain containing 3 | 0.0115 | No | 25.79 | 8 | 24.28 | 30.45 |
| ***AP2M1*** | adaptor related protein complex 2 mu 1 subunit | 0.0131 | No | 7.36 | 12 | 99.24 | 90.73 |
| ***CRIP1*** | cysteine rich protein 1 | 0.0202 | No | 20.16 | 5 | 113.86 | 111.57 |
| ***TMBIM6*** | transmembrane BAX inhibitor motif containing 6 | 0.0218 | No | 12.15 | 11 | 259.40 | 270.62 |
| ***RALB*** | RAS like proto-oncogene B | 0.0390 | No | 4.39 | 6 | 104.29 | 164.92 |
| ***RPL41*** | ribosomal protein L41 | 0.0477 | No | 6.92 | 6 | 467.15 | 479.03 |
| ***ITGAL*** | integrin subunit alpha L | 0.0494 | No | 7.78 | 11 | 142.95 | 152.56 |
| ***PTBP1*** | polypyrimidine tract binding protein 1 | 0.0494 | No | 13.19 | 9 | 48.16 | 52.49 |
| **Control vs. Anti-dsDNA+ENA+ SLE subset** | | | | | | | |
| ***FAM96A*** | family with sequence similarity 96 member A | 0.0068 | No | 15.11 | 5 | 36.69 | 41.05 |
| ***IFT20*** | intraflagellar transport 20 | 0.0079 | No | 23.36 | 6 | 17.39 | 19.29 |
| ***TIMM10*** | translocase of inner mitochondrial membrane 10 | 0.0108 | No | 21.65 | 3 | 18.67 | 56.34 |
| ***TNFAIP2*** | TNF alpha induced protein 2 | 0.0167 | No | 12.08 | 9 | 335.57 | 382.23 |
| ***AHCYL2*** | adenosylhomocysteinase like 2 | 0.0169 | Yes | 70.74 | 2 | 4.48 | 3.20 |
| ***HCK*** | HCK proto-oncogene, Src family tyrosine kinase | 0.0186 | No | 8.88 | 5 | 228.29 | 333.71 |
| ***ADIPOR1*** | adiponectin receptor 1 | 0.0431 | No | 5.31 | 3 | 213.16 | 190.58 |

***Abbreviations:*** *DIU, differential isoform usage. Total usage change reflects the aggregate shift in relative isoform proportions for a gene between the compared groups. “Major isoform switching” indicates whether the event was accompanied by a change in the dominant (most abundant) transcript isoform, versus a proportional change in isoform usage without a change in which isoform is dominant. Mean expression values are given in the units of the source quantification (e.g., normalized read counts). All genes listed passed the significance threshold of Q < 0.05.*
