## Supplementary table 4 for "Preferential Isoform Usage and Isoform Switching in Systemic Lupus Erythematosus Patients with Distinct Autoantibody Profile"

**Supplementary Table 4. Transcript- and protein-level features showing significant differential feature inclusion (DFI) in DIU gene across SLE patients’ subsets.**

| **Gene** | **Feature type** | **Feature ID(s)** | **Position** | **Ensembl transcript** | **Q value** | **Favoured condition** | **Isoform switch** | **ΔFI** |
| --- | --- | --- | --- | --- | --- | --- | --- | --- |
| **Anti-dsDNA+ subgroup** | | | | | | | | |
| ***GRAP2*** | miRNA binding site | hsa-miR-1207-5p | *Transcript* 39,971,050-39,971,058 | ENST00000460449 | 0.045 | Anti-dsDNA+ | No | 12.64 |
| ***ZSWIM8*** | miRNA binding site | hsa-miR-511-5p | *Transcript* 73,790,247-73,790,255 | ENST00000451629 | 0.010 | Control | No | 12.92 |
| ***DAZAP2*** | Protein domain | PF11029 | *Protein* 51,240,922-51,242,455 | ENST00000551313 | 2.14 × 10^-8^ | Anti-dsDNA+ | No | 7.19 |
|  | Compositional bias region | Pro-comp_bias | *Protein* 51,240,351-51,242,353 | ENST00000412716 | 9.91 × 10^-7^ | Control | No | 7.00 |
|  | miRNA binding sites (n = 10) | hsa-miR-106a-5p, hsa-miR-144-3p, hsa-miR-218-1-3p, hsa-miR-218-2-3p, hsa-miR-373-3p, hsa-miR-508-5p, hsa-miR-519e-3p, hsa-miR-520a-5p, hsa-miR-525-5p, hsa-miR-550a-5p | *Transcript* 51,242,504-51,242,747 | ENST00000552459 | 2.70 × 10^-5^ | Control | No | 6.94 |
|  | Simple repeat elements (n = 2) | Simple_repeat (×2) | *Transcript* 51,242,922-51,243,596 | ENST00000552459 | 9.55 × 10^-4^ | Control | No | 7.23 |
|  | miRNA binding sites (n = 22) | hsa-miR-124-3p, hsa-miR-1273f, hsa-miR-132-3p, hsa-miR-182-5p, hsa-miR-224-3p, hsa-miR-27a-3p, hsa-miR-27b-3p, hsa-miR-32-3p, hsa-miR-378a-3p, hsa-miR-378d, hsa-miR-422a, hsa-miR-4482-5p, hsa-miR-491-5p, hsa-miR-499a-3p, hsa-miR-506-3p, hsa-miR-5093, hsa-miR-522-3p, hsa-miR-7-5p, hsa-miR-767-5p, hsa-miR-891b, hsa-miR-92a-2-5p, hsa-miR-939-5p | *Transcript* 51,242,825-51,243,780 | ENST00000552459 | 0.005 | Control | No | 7.23 |
| ***TNFAIP2*** | Upstream open reading frame (uORF) | U0033 | *Transcript* 103,124,472-103,126,370 | ENST00000558056 | 0.002 | Control | No | 5.02 |
| **Anti-ENA+ subgroup** | | | | | | | | |
| ***IGHM*** | Poly(A) site | U0043 | *Transcript* 105,854,220-105,854,240 | ENST00000390559 | 8.26 × 10^-7^ | Anti-ENA+ | Yes | 67.42 |
|  | Poly(A) site | U0043 | *Transcript* 105,851,708-105,851,732 | ENST00000637539 | 8.26 × 10^-7^ | Control | Yes | 67.42 |
| ***CRIP1*** | Poly(A) site | U0043 | *Transcript* 105,488,767-105,488,791 | ENST00000392531 | 0.033 | Anti-ENA+ | Yes | 19.93 |
| ***BRI3*** | Repeat element | Simple_repeat | *Transcript* 98,281,748-98,281,793 | ENST00000297290 | 0.017 | Anti-ENA+ | Yes | 13.22 |
|  | Poly(A) site | U0043 | *Transcript* 98,291,503-98,291,524 | ENST00000297290 | 0.038 | Anti-ENA+ | Yes | 13.22 |
| ***FAM96A*** | Repeat | SINE/Alu | *Transcript* 64,073,788-64,073,817 | ENST00000558779 | 0.00511 | Control | No | 11.18 |
| **Anti-dsDNA+ENA+ subgroup** | | | | | | | | |
| ***TNFAIP2*** | Protein domain | PF06046 | *Protein* 103,127,627-103,135,342 | ENST00000560670 | 0.026 | Anti-dsDNA+ENA+ | No | 7.72 |
|  | Post-translational modification sites (n = 5) | Acetylation, Phosphoserine, Phosphothreonine, Phosphotyrosine, PTM_other | *Protein* 103,132,759-103,135,351 | ENST00000333007 | 0.044 | Anti-dsDNA+ENA+ | No | 7.72 |
|  | Upstream open reading frame (uORF) | U0033 | *Transcript* 103,124,472-103,126,370 | ENST00000558056 | 0.0105 | Control | No | 4.69 |
| ***ADIPOR1*** | 5’UTR motif | U0011 | *Transcript* 202,958,385-202,958,389 | ENST00000340990 | 0.0048 | Control | No | 5.31 |
| ***ADIPOR1*** | Upstream open reading frame (uORF) | U0033 | *Transcript* 202,951,099-202,958,232 | ENST00000340990 | 0.0415 | Control | No | 5.31 |

***Abbreviations:*** *DFI, differential feature inclusion; PAS, polyadenylation site; PTM, post-translational modification; uORF, upstream open reading frame; ΔFI, magnitude of change in feature-inclusion level between the compared groups. Coordinates are given in GRCh38 genomic numbering. “Favoured condition” indicates the group in which the feature was preferentially included. “Isoform switch” indicates whether the event was accompanied by a change in the dominant transcript isoform. All features listed passed the significance threshold of Q < 0.05.*

***Note:*** *Features mapping to the same gene, transcript isoform, position range, Q value and ΔFI (DAZAP2 in the anti-dsDNA+ subgroup; TNFAIP2 post-translational modification sites in the anti-dsDNA+ENA+ subgroup) have been consolidated into a single row listing all annotated Feature IDs (e.g., “n = 10”), reflecting one underlying isoform-usage event rather than independent findings. Full per-feature values are available in the source data.*
