## Supplementary table 5 for "Preferential Isoform Usage and Isoform Switching in Systemic Lupus Erythematosus Patients with Distinct Autoantibody Profile"

**Supplementary Table 5. Known interaction partners of *DAZAP2* and presence of the corresponding binding regions in the canonical and altered isoforms**

| **DAZAP2 domain / residues** | **Interacting protein (binding site)** | **Function / notes** | **Presence in canonical vs. altered isoform** | **Reference** |
| --- | --- | --- | --- | --- |
| – | **TCF-4**  (214–228 aa) | Knockdown of DAZAP2 reduces Wnt/β-catenin signaling | – | Lukas et al. 2009 |
| SH2 domains 1 and 3 | **IL17RB**  (329–347 aa) | SH2 domains 1 and 3 are essential for this interaction | **SH2 domain 1:** present in canonical; absent in altered isoform | Popova et al. 2012 |
| PPAY motif | **SMURF2** | – | Present in canonical; absent in altered isoform | Popova et al. 2012 |
| PPAY motif | **NEDD4** | – | Present in canonical; absent in altered isoform | Hamilton et al. 2001 |
| 67–168 aa | **eIF4GI**  (41–370 aa) | Overexpression of DAZAP2 reduces translation | Present in both isoforms | Kim et al. 2008 |

***Abbreviations:*** *aa, amino acids; SH2, Src homology 2; PPAY, proline-rich (PY-type) motif; TCF-4, transcription factor 4; IL17RB, interleukin-17 receptor B; SMURF2, SMAD-specific E3 ubiquitin protein ligase 2; NEDD4, neural precursor cell expressed developmentally down-regulated 4 (E3 ubiquitin ligase); eIF4GI, eukaryotic translation initiation factor 4G1. “–” indicates information not specified.*

***Note:*** *“Altered isoform” refers to the alternative DAZAP2 isoform showing changed usage in the differential isoform usage analysis.*
