## Supplementary figures and images for "Preferential Isoform Usage and Isoform Switching in Systemic Lupus Erythematosus Patients with Distinct Autoantibody Profile"

### Supplementary figure 1.tiff

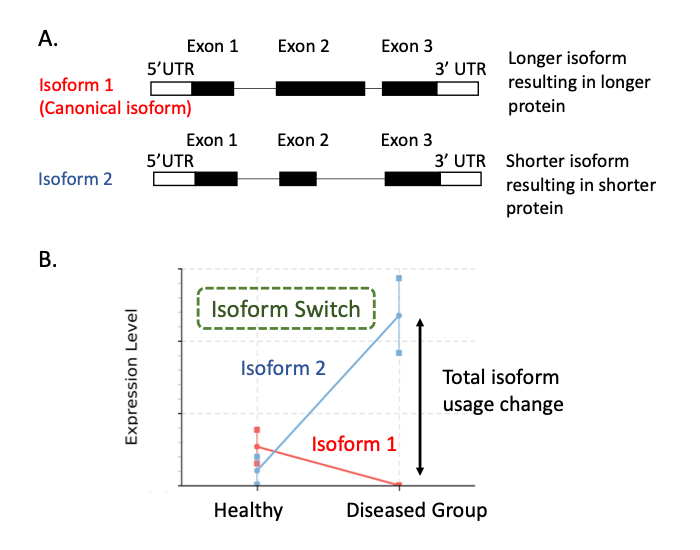

### Supplementary figure 2.tiff

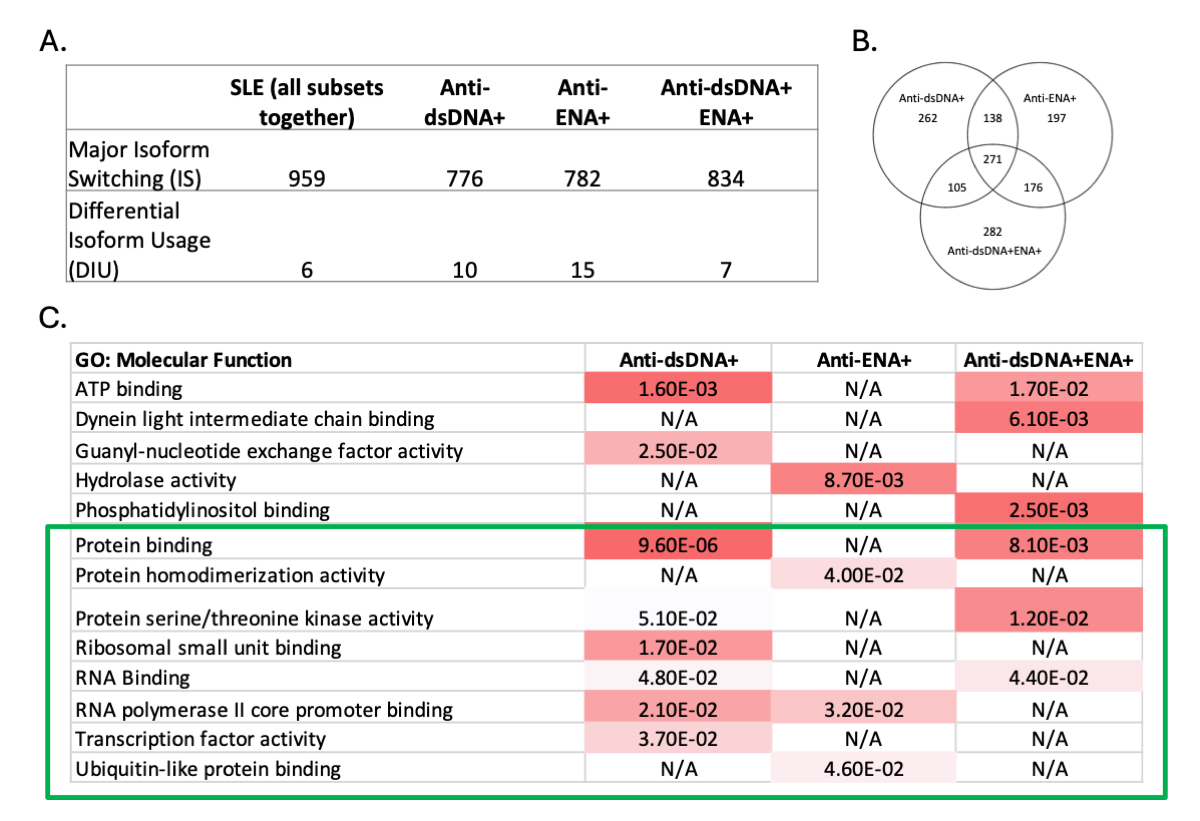

### Supplementary figure 3.tiff

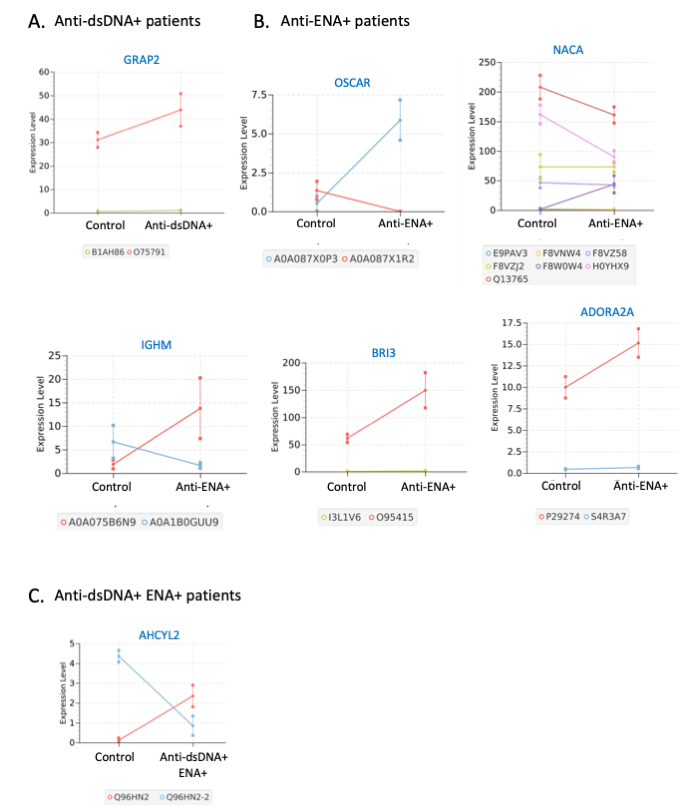

### Supplementary figure 4.tiff

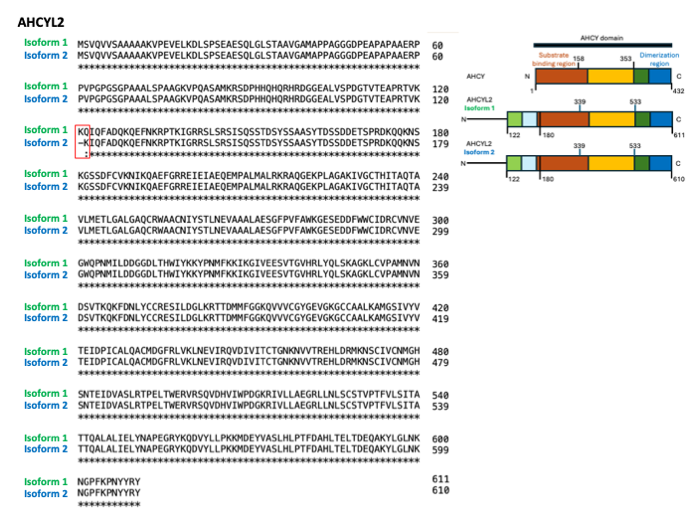

### Supplementary figure 5.tiff

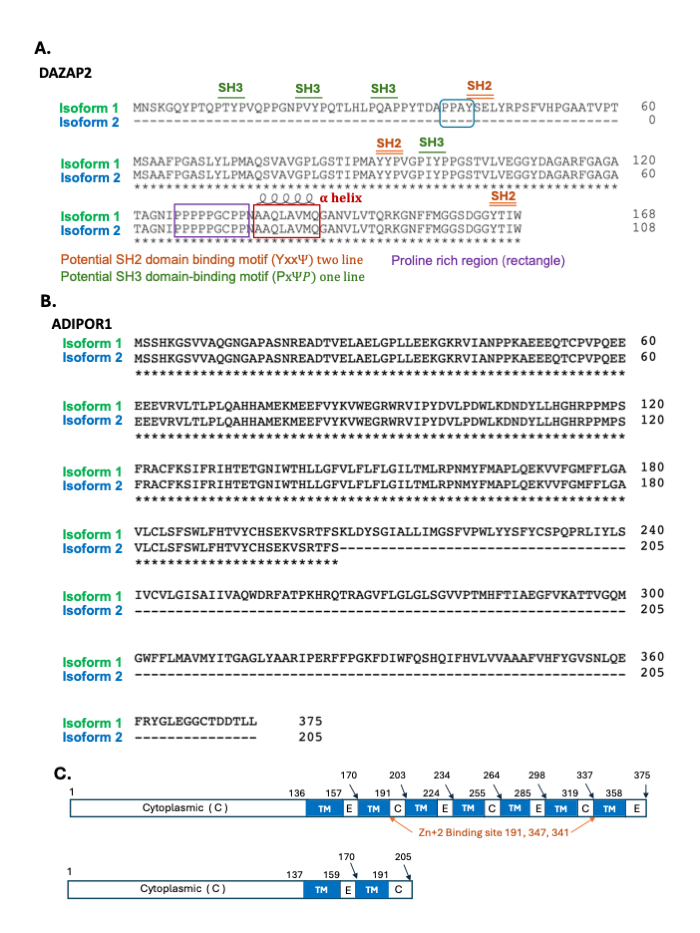
